# Stability and detection of nucleic acid from viruses and hosts in mosquito blood meals

**DOI:** 10.1101/2020.03.17.995126

**Authors:** Coyne Drummond, Mary E. Gebhardt, Maria Teresa Sáenz Robles, Giovanna Carpi, Isaiah Hoyer, Andrzej Pastusiak, Michael R. Reddy, Douglas E. Norris, James M. Pipas, Ethan K. Jackson

**Author notes:** Corresponding Author (MR).

## Abstract

Monitoring the presence and spread of pathogens in the environment is of critical importance. Rapid detection of infectious disease outbreaks and prediction of their spread can facilitate early responses of health agencies and reduce the severity of outbreaks. Current sampling methods are sorely limited by available personnel and throughput. For instance, xenosurveillance utilizes captured arthropod vectors, such as mosquitoes, as sampling tools to access blood from a wide variety of vertebrate hosts. Next generation sequencing (NGS) of nucleic acid from individual blooded mosquitoes can be used to identify mosquito and host species, and microorganisms including pathogens circulating within either host. However, there are practical challenges to collecting and processing mosquitoes for xenosurveillance, such as the rapid metabolization or decay of microorganisms within the mosquito midgut. This particularly affects pathogens that do not replicate in mosquitoes, preventing their detection by NGS or other methods. Accordingly, we performed a series of experiments to establish the windows of detection for DNA or RNA from human blood and/or viruses present in mosquito blood meals. Our results will contribute to trap design for mosquito-based xenosurveillance, including sample stabilization and ideal time spent from collection to NGS processing.

## Introduction

Mosquito-borne disease transmission represents a continued threat to human health and imposes an immense economic burden on at-risk populations (1–5). Monitoring human and zoonotic populations, especially in under-developed parts of the world, remains a challenge. Health organizations may lack the infrastructure necessary for adequate sampling and the reservoirs of future disease outbreaks are unknown. Detection of novel viruses for which we have no genetic data presents an additional challenge. The ability to rapidly detect emerging and established pathogens in humans and zoonotic reservoirs could allow for timely intervention and the potential to reduce outbreak severity by scaling up treatment production and prevention regimes, establishing quarantines and/or limiting contact with or culling afflicted animals. Recent studies have sampled wild mosquitoes to monitor the DNA and RNA of vertebrate blood and associated pathogens (6, 7). This technique, termed xenosurveillance, has been proposed for monitoring known and novel infectious diseases at (8–10).

The nucleic acid of both captured mosquitoes and blood meals can be analyzed by several approaches. PCR is an affordable method that has been used to screen for targeted hosts and pathogens in mosquito populations, using specific primer sequences for the taxa of interest (11). Recently described techniques utilize oligonucleotide hybridization to screen for panels of clinically important pathogens, especially viruses (12–14), but they are limited to known targets. Although these approaches are not ideal for the discovery of novel virus genomes without reference sequences, recent technological advances may allow detection of viruses presenting much greater sequence variability (15).

Next generation DNA and RNA sequencing (NGS), using platforms such as Illumina HiSeq and NovaSeq, are better suited for unbiased screening of all known and also novel virus sequences present in a particular sample. While some xenosurveillance studies have sequenced pools of mosquitoes (7), sequencing individual specimens offers the potential to link presence of a virus with a specific host reservoir. Unlike PCR and hybridization-based approaches, NGS combined with the proper analytical pipelines can be used not only to detect known viruses, but also to discover viruses with little or no sequence homology to reference sequences through *de novo* genome assembly (16).

There are practical challenges to storing captured mosquitoes, particularly blood fed mosquitoes, for metagenomic analysis. Once the specimens are collected, both DNA and RNA from the specimen, host and the blood meal are highly susceptible to degradation, especially in tropical and sub-tropical climates. Non-replicating viruses in particular do not escape the mosquito midgut and are rapidly digested. In order to improve specimen storage conditions and to determine the maximum time intervals between collection and molecular analytics, it is important to define the period during which nucleic acids can be successfully recovered and sequenced from blood meals. To this aim, we performed controlled blood feeds using *Aedes aegypti* Liverpool strain and *Anopheles stephensi* Liston mosquitoes. Viruses of varying capsid and genome types were added to blood meals, and the stability of viral genomes was monitored by real-time PCR and NGS to assess signal decay. We also tested the effects of cold storage on blood meal RNA and whether it favored nucleic acid preservation. Our experiments provide insights into optimal conditions for mosquito storage between collection and processing.

## Results

Previous xenosurveillance studies have identified viruses, including papillomaviruses (17), GB virus C, and hepatitis B virus (7), within blood meals of field-captured mosquitoes. We selected a panel of viruses that are commonly used in research laboratories and have diverse structural and genomic properties (Table 1) for our controlled feed experiments.

**Table 1.** Viruses used in blood feeds.

| Virus | Strain | Family | Genome Type | Genome Length | Enveloped | Stock Titer | Arbovirus |
| --- | --- | --- | --- | --- | --- | --- | --- |
| Human adenovirus | 5 | Adenoviridae | Linear dsDNA | 35.9 Kb | No | $2.1 \times 10^8$ | No |
| Sendai Virus | Cantell | Paramyxoviridae | Negative sense ssRNA | 15.4 Kb | Yes | $5 \times 10^7$ | No |
| Influenza | A | Orthomyxoviridae | Segmented negative sense ssRNA | 13.6 Kb | Yes | $1.5 \times 10^8$ | No |
| Dengue Virus 2 | C | Flaviviridae | Positive sense ssRNA | 10.7 Kb | Yes | 1*10<sup>6</sup> | Yes |

### Assessment of viral content by Real Time PCR (qPCR) in virus-fed mosquitoes

Nucleic acid content of mosquitoes fed with virus-containing blood meals was assessed by qPCR. Each mosquito blood meal was estimated to constitute an inoculum of 1,000 PFU (18). The first study included two RNA viruses, dengue virus 2 New Guinea strain C (DENV-2) and influenza A, which were fed to mosquitos either individually (5*10^5^ PFU / mL) or in combination (for a combined total of 5*10^5^ PFU / mL). Mosquitoes from the control group were either unfed or fed blood without virus.

DENV-2 is an arbovirus with tropism for *Ae. aegypti*, and was expected to be detectable by qPCR once it infected midgut tissues and started replicating with a 7 to 10-day extrinsic incubation period (19). In contrast, influenza A, a non-arbovirus and thus a proxy for xenosurveillance, was expected to have a much shorter window of detection, as the qPCR signal was expected to decay after the influenza A genome was digested or broken down. The combined feed was performed to determine whether the presence of more than one virus and/or an infected midgut might affect viral detection for xenosurveillance. 4 or 5 mosquitoes from each feed condition and time point were collected, and RNA was extracted for qPCR.

We found that influenza was readily detected by qPCR immediately after feeding and for the following 72 hours. However, there was little to no influenza signal at any time after four days post-feed (**Fig. 1A**). The influenza qPCR signal was otherwise unaffected by the presence of DENV-2 (**Fig. 1B**). The DENV-2 decay pattern was similar to that observed for influenza. A dramatic reduction of viral DENV-2 RNA signal was observed as early as 12 hours post feed, and little to no signal was apparent after 24 hours (**Fig. 1C**). As expected, the RNA levels began to increase between 100- and 200-hours post-feed, as the viral genome replicated in infected mosquito tissue (**Fig. 1D**).

**Fig. 1:**
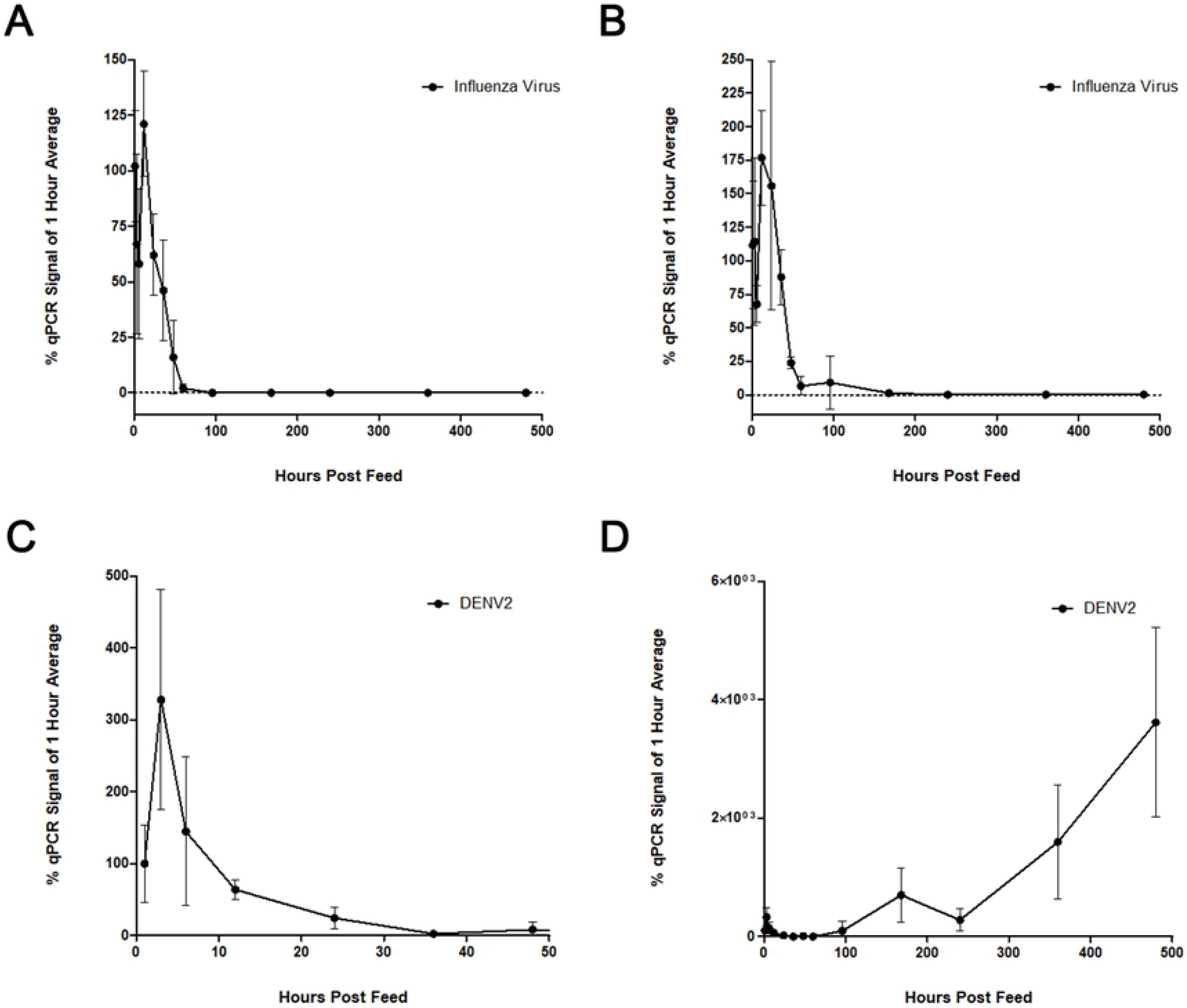
Viral blood feed time courses. *Ae. aegypti* were fed human blood spiked with 500 PFU of virus/ μL (approximately 1,000 PFU per 2 μL feed volume). For dual feeds this titration included 500 PFU of each virus. Post-feed levels of viral RNA were assessed by qPCR and normalized against endogenous mosquito *S7* (ribosomal protein S7) RNA levels. Each time point is represented in relation to the mean level at 1 hour post feed. (**A**) Influenza A levels following a single virus feed. (**B**) Influenza A levels after dual feed with DENV-2. (**C**) The first 40 hours of DENV-2 decay. (**D**) The full DENV-2 time course shows viral replication, beginning at approximately 100 hours, that occurs following the initial decay of the inoculum.

We next expanded the panel of viruses by incorporating a DNA virus, human adenovirus 5 (HAdV-5), and an RNA virus, Sendai virus (SeV), into separate *Ae. aegypti* blood feeds. Once again, viruses were titrated and added to fresh human blood (with 1.8 mg / mL EDTA added as preservative) to deliver 1,000 PFU to each mosquito, for an approximately 2 μL feed volume. In separate experiments, 14-16 mosquitoes per each time point and condition were allowed to feed until repletion over the course of one hour. They were then knocked down by chilling at 4°C for 5 minutes and subsequently sorted while cold and placed in a humidified chamber (27°C, 70RH, and 14/10-hour light/dark cycles) for the remainder of the time course. Approximately eight mosquitoes were removed for each collection time point and condition, killed by placing them at −20 °C and then transferred to −80 °C for storage. Both DNA and RNA were extracted from individual mosquitoes using column-based kits as described in the methods section. We assessed levels of human and viral DNA and RNA in the mosquito midguts via qPCR, using gene-specific primers targeting GAPDH or each separate virus (**Fig. 2**). We again found that signals for viral genomes of both SeV and HAdV-5 decayed appreciably 24 hours after feeding, and SeV was nearly undetectable by 48 hours. DNA and RNA from human cells within the blood, based on GAPDH levels, decayed somewhat more quickly, with a near complete loss of signal observed by 36 hours.

**Fig. 2:**
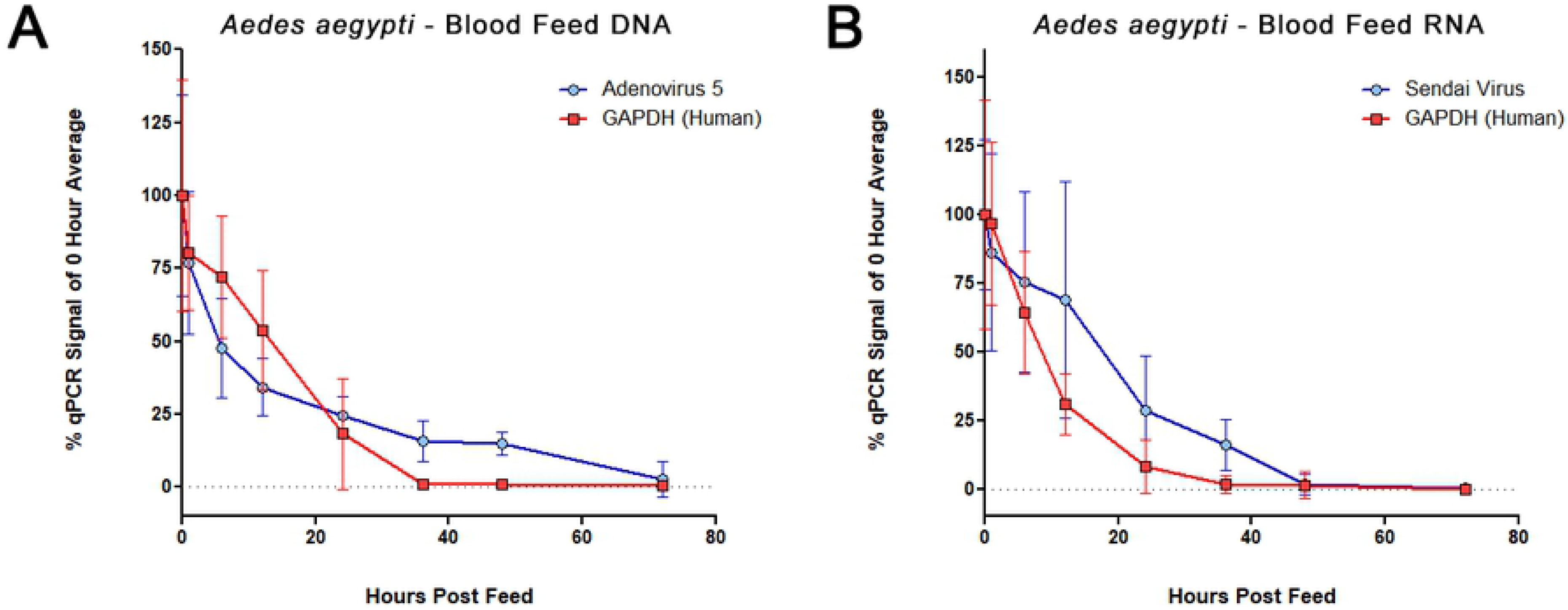
Mosquito blood feed qPCR time course. Real-time PCR for levels of DNA **(A)** and RNA **(B)** from a time course of blood fed *Ae. aegypti*, assessing the decay of human nucleic acid (GAPDH) and genomes of added viruses: HAdV-5 and SeV. Each point represents data from 14-16 individual mosquitoes, collected over two independent time courses. For each mosquito, virus and GAPDH signals were normalized to the mosquito endogenous *S17* (ribosomal protein S17) levels. Each time point is represented in relation to the mean level at feed time (time 0).

### Cold storage as a method for preservation of blood feed contents

The previous experiments suggest blood meal contents can be detected for up to 36 to 48 hours in mosquitoes that fed immediately before capture. However, a mosquito may enter traps many hours after feeding and after blood meal digestion has started. Blood meal RNA in particular maybe at particular risk of decaying due to high ambient temperatures, particularly in the heat of tropical and subtropical climates. Therefore, it would be of interest to better preserve nucleic acid directly at the point of capture to slow the digestion of the mosquito blood meal. We thus tested the possible preservative effects of cold storage (4°C) on human SeV RNA within mosquito blood meals. As described for previous experiments, *Ae. aegypti* were fed human blood containing 5*10^5^ PFU/mL SeV.

After 12 hours in the insectary incubation chamber, cups of live fed mosquitoes were either moved to a 4°C chamber for 24 hours or left at 27°C for the same period. All mosquitoes held at 4°C were dead after 24 hours. Abdomen sizes of 4°C-held mosquitoes more closely resembled those of 12-hour mosquitoes than did those from mosquitoes that were kept live at 27°C for the entire 36 hours (not measured/shown). All specimens were chilled at −20°C for 10 minutes and stored at −80°C until processed. RNA was extracted and assessed by real-time PCR. Levels of human GAPDH and SeV RNA were found to be significantly higher in mosquitoes that were held for 24 hours at 4°C (p=.0003), suggesting that the blood meal and contents were better preserved by cold temperature (**Fig. 3**). There was no significant difference between the RNA levels of either GAPDH (p=0.2403) or SeV (p=0.0758) recovered from mosquitoes stored from 12 to 36 hours at 4°C and from those directly collected after 12 hours at 27°C. We performed additional tests with bovine blood meals (**Fig. 4**) and found higher levels of host-blood DNA and RNA in the samples stored at cold temperature. These results indicate that blood meal nucleic acid content can be preserved via temperature control. However, the data do not address whether the signal decay that occurs at warmer temperatures is primarily due to the digestive activity of the mosquito or to the inherent instability of the DNA and RNA.

**Fig. 3.**
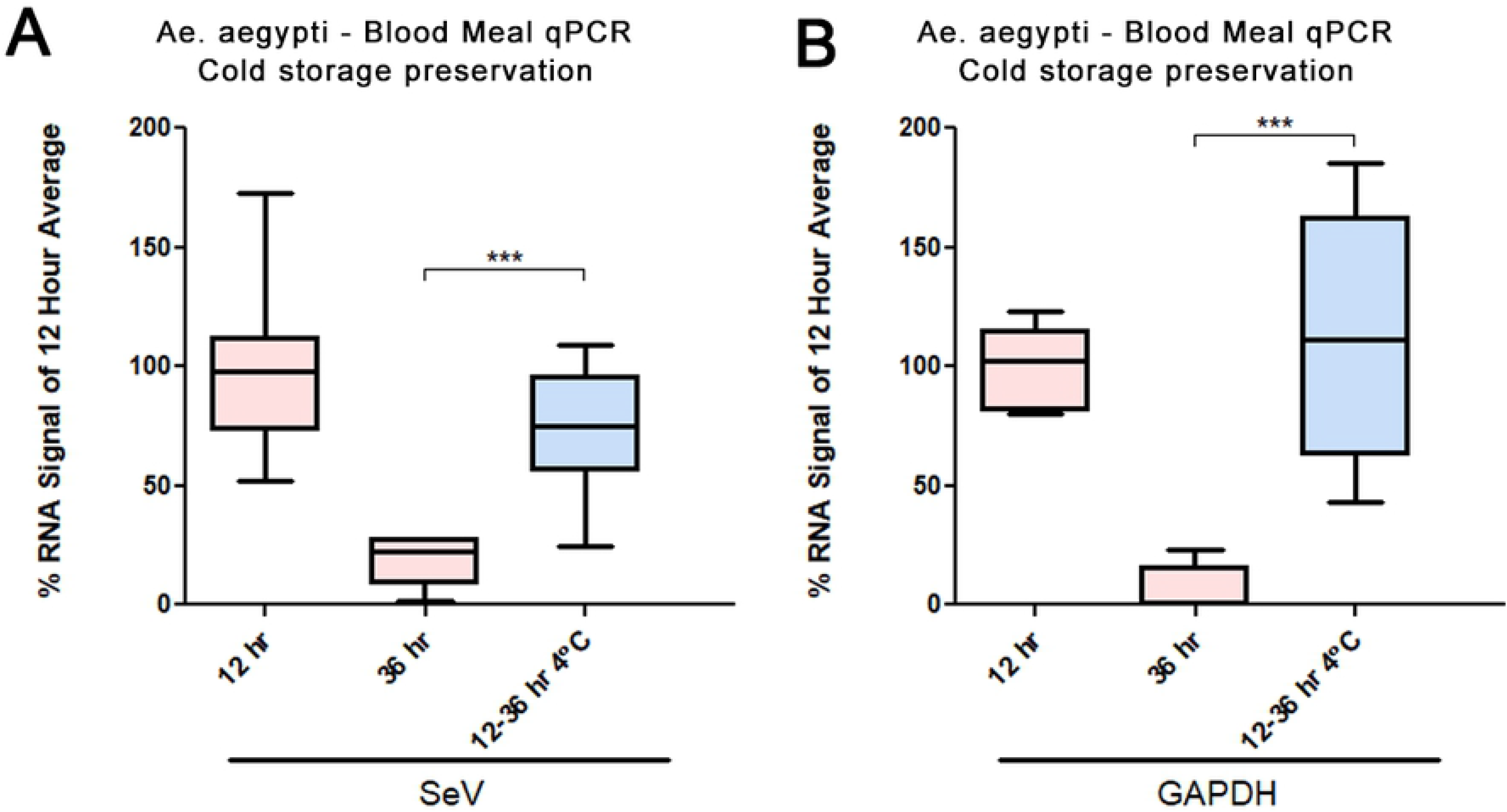
Cold preservation for blood meal RNA. RNA levels of SeV and GAPDH in blood meals at 12- and 36-hours post feed, either from mosquitoes held at regular insectary temperatures (27°C) and conditions, or from samples held for 24 hours (12 to 36 hours post-feed) at 4°C. CTs are normalized to the corresponding values for *Ae. aegypti* S17 and are shown as percent of mean at 12-hour signal. No significance was observed between samples kept at either temperature for 12 hour by application of two-tailed Welch’s t-test, but there was a significant reduction in the RNA signals from samples kept at 27°C for 36 hours (p=.0003 ***). Eight samples were analyzed for each time point, except those processed after 12 hours (seven samples).

**Fig. 4.**
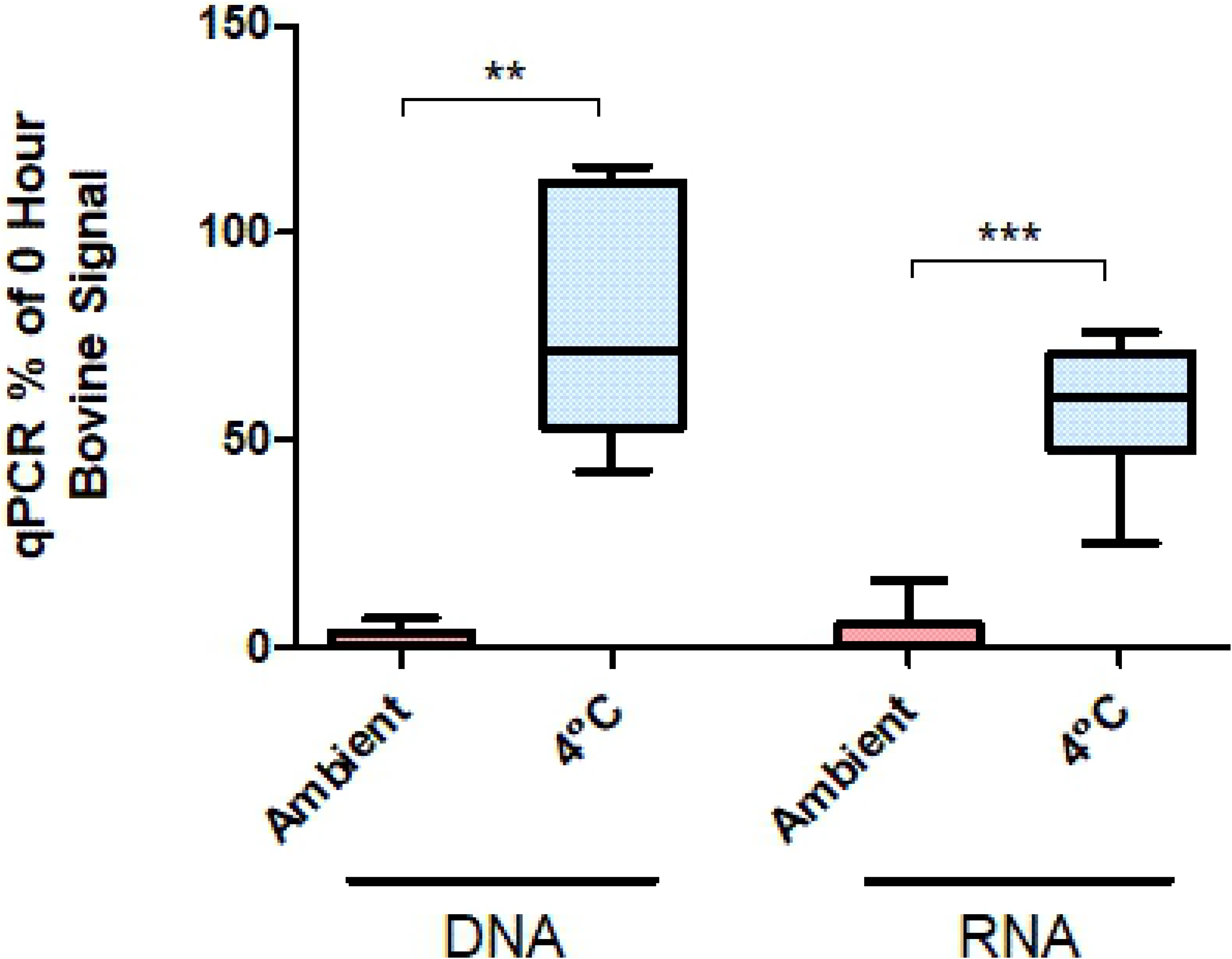
Bovine blood nucleic acid stability in mosquito midguts. *An. stephensi* were held at 21° C (RT) or 4°C for 48 hours immediately post-feeding, or frozen immediately after feeding (−80°C). DNA and RNA were isolated from individual mosquitoes and bovine nucleic acid levels were determined with qPCR using primers targeting bovine cytochrome B. Results are shown as percentages relative to the mean of samples frozen immediately after feeding. ** − p = 0.024 (n =5). *** − p < .0001 (n=7).

## Methods

### Virus production and titration

Previous xenosurveillance studies have identified viruses, including papillomaviruses (17), GB virus C, and hepatitis B virus (7), within blood meals of field-captured mosquitoes. We selected a panel of viruses that are commonly used in research laboratories and have diverse structural and genomic properties (**Table 1**) for our controlled feeds experiments. Prior to use, all virus stocks were suspended in PBS and titrated by plaque assay to determine concentrations in plaque forming units (PFU). For each feeding experiment, 5*10^5^ PFU of virus was added to 1.0 mL human blood. With this prepared blood meal and estimating that each female mosquito would imbibe 2 μL (18), each blood meal would constitute an inoculum of 1,000 viral PFU.

#### SeV

The SeV Cantell strain stock of 1.6*10^4^ HAU/mL was used to create working stocks of 800 HAU/mL. Viruses were titrated on Vero cells with a modified plaque assay technique, based on pre-existing protocols (20, 21), to determine the titers in plaque forming units (PFU). Briefly, viral stock dilutions were plated onto confluent Vero cells in serum-free Dulbecco’s minimal essential medium. Viruses were allowed to bind cells for 1 hour at 37°C. Cells were washed twice with PBS, and then MEM overlay medium containing 1.2 μg/mL trypsin and 1% agarose was applied. After 72 hours, agarose plugs were removed, and monolayers were stained with crystal violet. The original (1.6*10^4^ HAU/mL) and 800 HAU/mL stocks contained 10^9^ and 5*10^7^ PFU/mL, respectively, and the later was used for all mosquito blood feed experiments.

#### HAdV-5

Stocks were prepared by inoculating 10 cm plates of confluent HEK-293T cells with 10^7^ PFU HAdV-5. After 4 days, most cells were detached from the plates. Cells were scraped and media was collected. After three successive freeze/thaw cycles, cell debris was pelleted at 5,000 RPM, and the supernatant was used for viral stocks. HAdv-5 titers were determined via plaque assay on Vero cells, using a 0.5% agarose overlay. A stock of 2.1*10^8^ PFU/mL was used for all subsequent feed experiments.

#### DENV-2 and influenza

Dengue 2 (New Guinea C) and influenza A viral stocks each contained 1*10^6^ PFU / mL (the influenza working stock was diluted from an original stock at 1.5*10^8^ PFU / mL with DMEM).

### Mosquito rearing

*Aedes aegypti* Liverpool strain and *Anopheles stephensi* Indian wild type strain were used for the feeding experiments. *Ae. aegypti* were reared in the Johns Hopkins Malaria Research Institute Insectary at the Bloomberg School of Public Health under standard conditions. *An. stephensi* were purchased from the Center for Global Infectious Disease Research in Seattle. Adult mosquitoes were maintained at 27°C and 70RH, with access to 10% sucrose solution *ad libitum*. All female mosquitoes used in the feeding experiments were 3-8 days post-eclosion and were starved for 4-12 hours prior to blood feeding.

### Mosquito blood feeds

Bovine blood feeding experiments were performed with *An. stephensi* at Microsoft Research. Whole heparinized bovine blood (Hemostat, Dixon, CA) was purchased and after starving for 12 hours mosquitoes were fed via an artificial membrane feeder (Hemotek, Blackburn, UK). After feeding 7 to 14 fully engorged mosquitoes were transferred to each screened 50 mL tube. Tubes were held at 21° C or 4° C for 48 hours. At 48 hours mosquitoes were moved to −80°C until processed.

All virus feeding experiments were performed with *Ae. aegypti* under containment conditions. Twelve hours prior to blood feeds, female mosquitoes were identified and removed from rearing cages. Approximately 60 to 80 females were aspirated into each feeding cup. Fresh whole human blood, preserved with 1.8 mg/mL EDTA, was supplied by the Johns Hopkins University Parasite Core within two days prior to feeding experiments. Blood was allowed to warm to room temperature and virus stock was added such that the final concentration of virus was 5*10^5^ PFU/mL, or 1,000 PFU per each expected 2 μL blood meal. Parafilm-covered glass feeders were placed on each feeding cage and warmed to 37°C. Cages of unfed ‘control’ mosquitoes were kept in the incubation chamber throughout the feed. Each feeder was loaded with 300 μL of blood, with or without virus, and mosquitoes were allowed to feed for up to 60 minutes before being knocked down and visibly fed mosquitoes sorted into cups at 4°C. The 0-hour time point was recorded and collected when mosquito cups were returned to 27°C incubation chambers. At each time point, one cup with 8-16 mosquitoes per condition was placed at −20°C for 10 minutes. Killed mosquitoes were sorted into individual 1.5mL microfuge tubes and stored at −80°C.

### Evaluating cold storage

As described for previous experiments, *Ae. aegypti* were fed human blood containing 5*10^5^ PFU/mL SeV. After 12 hours in the insectary environmental chamber (27°C), cups of live fed mosquitoes were either moved to a 4°C chamber for 24 hours or left at 27°C for the same period. All specimens were chilled at −20°C for 10 minutes and transferred individually to 0.6 mL microfuge tubes and stored at −80°C until processed.

### Extraction and quantitative PCR

Whole mosquitoes were stored at −80° for up to two months until shipped to the University of Pittsburgh for nucleic acid extraction and analysis. Mosquito tissues were disrupted with a FastPrep-24 benchtop homogenizer (MP Biomedical, Irvine, CA), using two sixty second pulses at 4.0 m/s. Total DNA and RNA were extracted separately from each individual mosquito using AllPrep DNA/RNA (Qiagen, Hilden, Germany), according to the manufacturer’s instructions. Relative amounts of mosquito, human, and viral DNA (S17, S7, GAPDH, and HAdV-5) and RNA (S17, S7, GAPDH, SeV, DENV-2, and influenza) were assessed by real-time PCR with Power SYBR 1-Step Kit, using the AB 7900HT (Applied BioSystems, Foster City, CA) real-time PCR machine. Gene-specific primers sets used included human GAPDH (F: 5-’AATCCCATCACCATCTTCCAGG-3’ / R: 5’-GCCTCCCCAAAGCACATTTC-3’), *Ae. aegypti* ribosomal protein S17 (F: 5’-CACTCCCAGGTCCGTGGTAT-3’ / R: 5’-GGACACTTCCGGCACGTAGT-3’), *An. stephensi* ribosomal protein S7 (F: 5’-GGTGCACCTGGATAAGAACCA-3’ / R: 5’-CGGCCAGTCAGCTTCTTGTAC-3’), Dengue (F: 5’-AGGACYAGAGGTTAGAGGAGA-3’ / R: 5’-CGYTCTGTGCCTGGAWTGAT-3’), influenza (F: 5’-GGGTTTGTGTTCACGCTCAC-3’ / R: 5’-GGCATTTTGGACAAAGCGTCTAC-3’) SeV (F: 5’-CAGAGGAGCACAGTCTCAGTGTTC-3’ / R: 5’-TCTCTGAGAGTGCTGCTTATCTGTGT-3’), HAdV-5 (F: 5’-TTGTGGTTCTTGCAGATATGGC-3’ / R: 5’-TCGGAATCCCGGCACC-3’), and bovine cytochrome B (F: 5’-CGGAGTAATCCTTCTGCTCACAGT-3’ / R: 5’-GGATTGCTGATAAGAGGTTGGTG-3’). Real-time signal of blood feed nucleic acid was normalized to the signal of mosquito ribosomal protein S7 or S17 (for *Anopheles* and *Aedes*, respectively).

## Discussion

The use of xenosurveillance with blood-feeding arthropods presents an enticing possibility for sampling a wide variety of animal and human blood in order to monitor for viruses and other pathogens. High-throughput processing of mosquitoes, collected from strategically placed traps, could access a much broader range of hosts species, with wider geographic reach and at potentially much lower cost than traditional methods. Here, we performed a series of proof-of-principle experiments designed to detect known, non-replicating viruses in mosquito blood meals. These experiments were meant to replicate the transient presence of non-arboviruses being passed along from host to mosquito midgut via the blood meal.

The breakdown of nucleic acids and the digestive activity in mosquito midguts limits the potential window of detection for blood meals. Our blood feed time courses confirmed that DNA and RNA (human, bovine, Influenza, Adenovirus, and Sendai virus) present in blood meals decayed within several days of feeding, while mosquito nucleic acid levels remained constant (**Fig 1, 2, 4**). Interestingly, detection of viral genomes appeared to decay more slowly than that of human blood (**Fig 2**), perhaps as a consequence of the encapsidated viral genomes being better protected than cellular DNA and RNA. However, this difference could also be due to relative transcript/genome abundance, and NGS would be a better platform to assess this effect.

The optimal 24- to 36- hour window for detection of host blood and viral nucleic acid suggests that mosquito traps without any preservative capabilities should be collected, at minimum, daily. Given that mosquitoes will not necessarily enter the trap immediately after feeding, more regular trap collection intervals are probably advisable. Alternatively, we have found that blood meal detection can be extended to a longer period post-feeding (from 12 to 36 hours [**Fig. 3**] and 0 to 48 hours [**Fig. 4**]) with minimal signal loss by qPCR. Additional experiments are needed to determine the ideal temperature for preservation, and to determine what would be feasible for a remotely deployed trap.

## Conclusions

Screening the DNA and RNA of blood-feeding insects may be beneficial for monitoring the presence and spread of pathogens circulating within their vertebrate blood meal hosts. This technique, termed xenosurveillance, may also provide insights into the emergence of novel viruses and the animal and human reservoirs of known viruses. Blooded mosquitoes collected for xenosurveillance must be collected shortly after feeding, and the blood meal and its nucleic acid content must either be processed or preserved before decaying or digestion. Here, we establish the time constraints for detecting selected DNA and RNA virus genomes in mosquito blood meals and assess the effects of cold treatment to stabilize the samples and help preserve these genomes.

## Acknowledgements

HAdV-5 and Influenza stocks were kindly provided by Dr. Gary Ketner and Dr. Andrew Pekosz, respectively, at Johns Hopkins University Bloomberg School of Public Health. We thank Dr. Jenny Carlson and Hannah MacLeod for assistance with the mosquito feeds. DEN and MEG were supported in part by the Hopkins Malaria Research Institute and Bloomberg Philanthropies, and MEG also received support from NIH 5T32AI007417.

## References

1. Sachs J, Malaney P. The economic and social burden of malaria. Nature. 2002;415(6872):680–5.

2. Barrett ADT. Economic burden of West Nile virus in the United States. Am J Trop Med Hyg. 2014;90(3):389–90.

3. Castro MC, Wilson ME, Bloom DE. Disease and economic burdens of dengue. Lancet Infect Dis. 2017;17(3):e70–e8.

4. Lee BY, Alfaro-Murillo JA, Parpia AS, Asti L, Wedlock PT, Hotez PJ, et al. The potential economic burden of Zika in the continental United States. PLoS Negl Trop Dis. 2017;11(4):e0005531.

5. Alfaro-Murillo JA, Parpia AS, Fitzpatrick MC, Tamagnan JA, Medlock J, Ndeffo-Mbah ML, et al. A cost-effectiveness tool for informing policies on Zika virus control. PLoS Negl Trop Dis. 2016;10(5):e0004743.

6. Kading RC, Biggerstaff BJ, Young G, Komar N. Mosquitoes used to draw blood for arbovirus viremia determinations in small vertebrates. PLoS One. 2014;9(6):e99342.

7. Fauver JR, Weger-Lucarelli J, Fakoli LSI, Bolay K, Bolay FK, Diclaro JWI, et al. Xenosurveillance reflects traditional sampling techniques for the identification of human pathogens: A comparative study in West Africa. PLoS Negl Trop Dis. 2018;12(3):e0006348.

8. Brinkmann A, Nitsche A, Kohl C. Viral metagenomics on blood-feeding arthropods as a tool for human disease surveillance. Int J Mol Sci. 2016;17(10):E1743.

9. Fauver JR, Gendernalik A, Weger-Lucarelli J, Grubaugh ND, Brackney DE, Foy BD, et al. The Use of Xenosurveillance to Detect Human Bacteria, Parasites, and Viruses in Mosquito Bloodmeals. Am J Trop Med Hyg. 2017;97(2):324–9.

10. Grubaugh ND, Sharma S, Krajacich BJ, Fakoli Iii LS, Bolay FK, Diclaro Ii JW, et al. Xenosurveillance: A Novel Mosquito-Based Approach for Examining the Human-Pathogen Landscape. PLoS Negl Trop Dis. 2015;9(3):e0003628.

11. Brugman VA, Hernandez-Triana LM, Prosser SW, Weland C, Westcott DG, Fooks AR, et al. Molecular species identification, host preference and detection of myxoma virus in the Anopheles maculipennis complex (Diptera: Culicidae) in southern England, UK. Parasit Vectors. 2015;8:421.

12. Deviatkin AA, Lukashev AN, Markelov MM, Gmyl LV, Shipulin GA. Enrichment of viral nucleic acids by solution hybrid selection with genus specific oligonucleotides. Sci Rep. 2017;7(1):9752.

13. Briese T, Kapoor A, Mishra N, Jain K, Kumar A, Jabado OJ, et al. Virome capture sequencing enables sensitive viral diagnosis and comprehensive virome analysis. MBio. 2015;6(5):e01491–15.

14. Wylie TN, Wylie KM, Herter BN, Storch GA. Enhanced virome sequencing using targeted sequence capture. Genome Res. 2015;25(12):1910–20.

15. Metsky HC, Siddle KJ, Gladden-Young A, Qu J, Yang DK, Brehio P, et al. Capturing sequence diversity in metagenomes with comprehensive and scalable probe design. Nat Biotechnol. 2019;37(2):160–8.

16. Cantalupo PG, Pipas JM. Detecting viral sequences in NGS data. Curr Opin Virol. 2019;39:41–8.

17. Ng TF, Willner DL, Lim YW, Schmieder R, Chau B, Nilsson C, et al. Broad surveys of DNA viral diversity obtained through viral metagenomics of mosquitoes. PLoS One. 2011;6(6):e20579.

18. Ogunrinade A. The measurement of blood meal size in Aedes aegypti (L.). Afr J Med Med Sci. 1980;9(1-2):69–71.

19. Salazar MI, Richardson JH, Sanchez-Vargas I, Olson KE, Beaty BJ. Dengue virus type 2: replication and tropisms in orally infected Aedes aegypti mosquitoes. BMC Microbiol. 2007;7:9.

20. Buggele WA, Horvath CM. MicroRNA profiling of Sendai virus-infected A549 cells identifies miR-203 as an interferon-inducible regulator of IFIT1/ISG56. J Virol. 2013;87(16):9260–70.

21. Zimmermann M, Armeanu-Ebinger S, Bossow S, Lampe J, Smirnow I, Schenk A, et al. Attenuated and protease-profile modified sendai virus vectors as a new tool for virotherapy of solid tumors. PLoS One. 2014;9(3):e90508.

